# tVTA controls dual dopaminergic inputs to the external Globus Pallidus

**DOI:** 10.64898/2026.04.20.719622

**Authors:** Margaux Lebouc, Giulia R. Fois, Alessandro Bilella, Jérôme Baufreton, Michel Barrot, François Georges

## Abstract

Midbrain dopamine (DA) neurons critically regulate basal ganglia function through their widespread projections. While the nigrostriatal pathway is well characterized and represents the dominant source of DA in the basal ganglia, other nuclei such as the external Globus Pallidus (GPe) also receive dopaminergic innervation, yet no consensus exists about its precise anatomical origin. In addition, the GABAergic tail of the ventral tegmental area (tVTA) provides a major inhibitory input to midbrain DA neurons, but its influence over DA pathways to the GPe remains unknown. In the rat, we combined retrograde tracing, immunohistochemistry, and ex vivo electrophysiology to identify distinct populations of DA neurons in the substantia nigra pars compacta (SNc) and ventral tegmental area (VTA) that project to the GPe and display distinct electrophysiological properties. Using optogenetics and electrophysiology, we also demonstrate that these GPe-projecting DA neurons receive powerful inhibitory input from the tVTA. Together, our findings define both the origin and inhibitory control of dopaminergic innervation to the GPe, revealing a previously unrecognized disynaptic circuit (tVTA→DA→GPe) that refines our understanding of basal ganglia circuit function.

## INTRODUCTION

The basal ganglia (BG) are a network of interconnected structures crucial for the selection and initiation of appropriate motor behavior (Alexander and Crutcher, 1990; Graybiel et al., 1994). Traditionally, the external globus pallidus (GPe) was considered as a relay nucleus within the indirect pathway, receiving inputs primarily from the striatum and projecting to the subthalamic nucleus (STN) and basal ganglia output structures, namely the substantia nigra pars reticulata (SNr) and the internal globus pallidus (GPi) (Albin et al., 1989; Gerfen et al., 1990). However, accumulating evidence has revealed the cellular and functional heterogeneity of the GPe, highlighting its ability to integrate and respond to inputs from multiple brain regions, including the cortex, thalamus, and pedunculopontine nucleus (Abdi et al., 2015; Aristieta et al., 2021; Dong et al., 2021; Karube et al., 2019; Ma et al., 2025; Mallet et al., 2016, 2012).

Dopaminergic modulation is a central feature of BG function. Dopamine (DA) neurons from the substantia nigra pars compacta (SNc) predominantly innervate the striatum, where they regulate motor output through modulation of the direct and indirect pathways. In contrast, DA neurons from the ventral tegmental area (VTA) primarily project to limbic and cortical regions, including the nucleus accumbens (NAc) and prefrontal cortex (PFC), where they contribute to learning and motivational behaviors (Sesack and Grace, 2010). Although less dense than striatal innervation, dopaminergic projections to the GPe have been described (Debeir et al., 2005; Parent et al., 1990), and are thought to modulate pallidal neuron activity (Eid and Parent, 2015). Indeed, intrapallidal dopamine application increases GPe neuronal firing whereas dopamine receptor antagonists (D1 or D2) reduce firing and impair motor function (Mamad et al., 2015; Querejeta et al., 2001). Furthermore, hotspots of DA release in the GPe have been reported supporting a patchy DA modulation in the GPe (Meszaros et al., 2018). However, the precise anatomical origin of these dopaminergic inputs remains uncertain. Previous anatomical studies have primarily focused on nigro-pallidal projections (Eid and Parent, 2015), but tracing studies in non-human primates have also identified the VTA as a potential source of dopaminergic input to the GPe (Charara and Parent, 1994), suggesting a possible dual origin. Based on these observations, we hypothesized that the GPe receives dopaminergic inputs from both SNc and VTA neurons.

The tail of the ventral tegmental area (tVTA), also named the rostromedial tegmental nucleus (RMTg), is a major GABAergic structure that exerts powerful inhibitory control over midbrain DA neurons (Jalabert et al., 2011; Jhou, 2021; Jhou et al., 2009a, 2009b; Kaufling et al., 2010, 2009; Lecca et al., 2012; Matsui and Williams, 2011). Optogenetic activation of tVTA afferents induces GABA_A_ receptor-mediated inhibitory postsynaptic currents in both SNc and VTA DA neurons (Matsui and Williams, 2011), and tVTA inhibition or ablation has been shown to influence motor performance and learning (Bourdy et al., 2014; Faivre et al., 2020). These findings raise the possibility that tVTA may also regulate dopaminergic projections to the GPe.

Here, we tested the hypothesis that tVTA exerts inhibitory control over GPe-projecting dopaminergic neurons. To this end, we combined retrograde tracing from the GPe with immunohistochemistry, *ex vivo* electrophysiology, and optogenetics to characterize the anatomical origin, intrinsic properties, and synaptic regulation of this previously understudied dopaminergic pathway.

## RESULTS

### Dopaminergic innervation of the GPe originates from both the SNc and VTA

To identify the origin of dopaminergic inputs to the GPe, we injected Fluoro-Gold, a retrograde tracer, into this structure (Figure 1A). Retrograde labeling revealed a substantial number of neurons in both the SNc and the VTA (Figure 1B). Among retrogradely labeled neurons, 90% in the SNc and 70% in the VTA were tyrosine hydroxylase (TH)-positive and therefore dopaminergic (Figure 1C). This difference is not surprising, since the VTA is known for its high cellular heterogeneity (Sesack and Grace, 2010). These results indicate that, following Fluoro-Gold injection, approximately 40% of the dopaminergic innervation of the GPe originates from the SNc and 60% from the VTA, supporting the conclusion that both regions contribute substantially to GPe dopaminergic input (Figure 1D).

**Figure 1:**
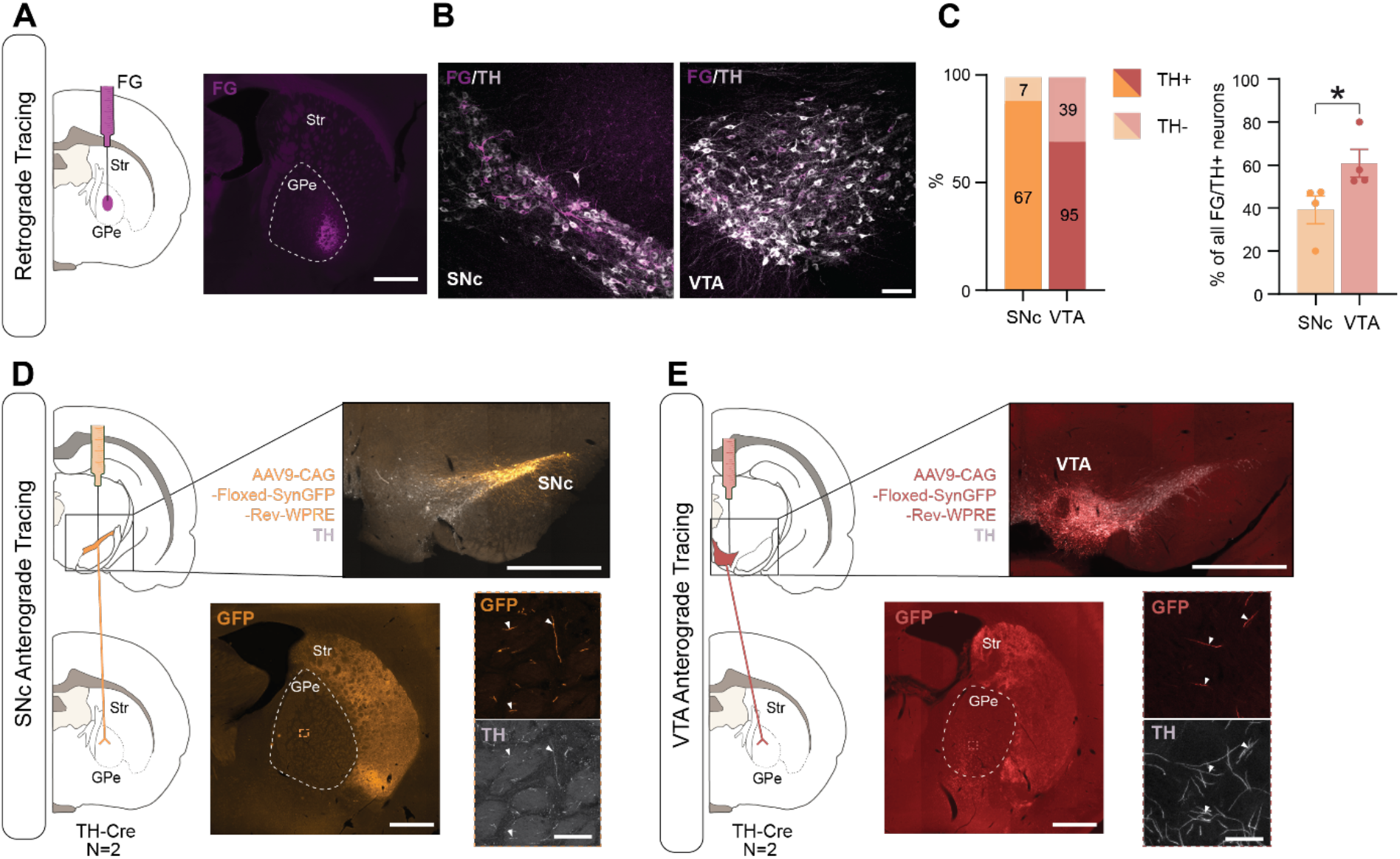
The GPe receives a dopaminergic innervation from both the SNc and the VTA. **A**. Retrograde injection of Fluoro-Gold (FG) into the GPe. **B**. Left, FG-positive (FG^+^) and TH-positive (TH^+^) neurons in the SNc. Right, FG^+^ and TH^+^ neurons in the VTA. **C**. Left, proportion of TH^+^ (dark) and TH^-^ (light) neurons among FG^+^ cells in SNc (orange) and VTA (red). The number depicted on each bar indicates the total number of cells counted. Right, percentage of FG^+^/TH^+^ neurons in SNc and VTA. **D**. Anterograde labeling of SNc dopaminergic fibers. Top, Injection site in SNc. Bottom, SynGFP^+^ fibers in GPe and STR and high magnification view of fibers in GPe. **E**. Anterograde labeling of VTA dopaminergic fibers. Top, Injection site in VTA. Bottom, SynGFP^+^ fibers into the GPe and STR and high magnification view in GPe. Scale bars: 1 mm, 100 μm and 50 μm.

The GPe contains numerous passage fibers, including dopaminergic axons of the nigrostriatal pathway (Eid and Parent, 2015). To confirm that retrograde labeling reflected true synaptic inputs rather than uptake by passing fibers, we used an anterograde approach to label dopaminergic synaptic terminals in the GPe. TH-Cre rats were injected with a Cre-dependent viral vector encoding synaptophysin-GFP (Syn-GFP). Injections were confined to either the SNc or the VTA (Figure 1D-E). Although the signal in the GPe was weaker than in the striatum, numerous Syn-GFP-positive fibers were observed at high magnification. All Syn-GFP-labeled fibers were TH-positive, confirming their dopaminergic identity. Some TH-positive fibers lacked Syn-GFP labeling, indicating they originated from the alternative dopaminergic source (VTA for SNc injection, SNc for VTA injection). These results corroborate the retrograde tracing data, demonstrating that dopaminergic synaptic terminals in the GPe arise from both the SNc and VTA.

### Distinct electrophysiological properties of SNc- and VTA-derived GPe-projecting DA neurons

DA neurons exhibit well-characterized electrophysiological properties, which can differ depending on their projection target (Ford et al., 2006; Lammel et al., 2008). To examine the properties of GPe-projecting SNc and VTA neurons, we performed patch-clamp recordings of retrogradely labeled cells (Figure 2). Multiple retrograde tracers were used, including FG, CTb, and retrobeads. Post-hoc immunostaining confirmed the dopaminergic identity of recorded neurons by TH expression (Figure 2B).

**Figure 2.**
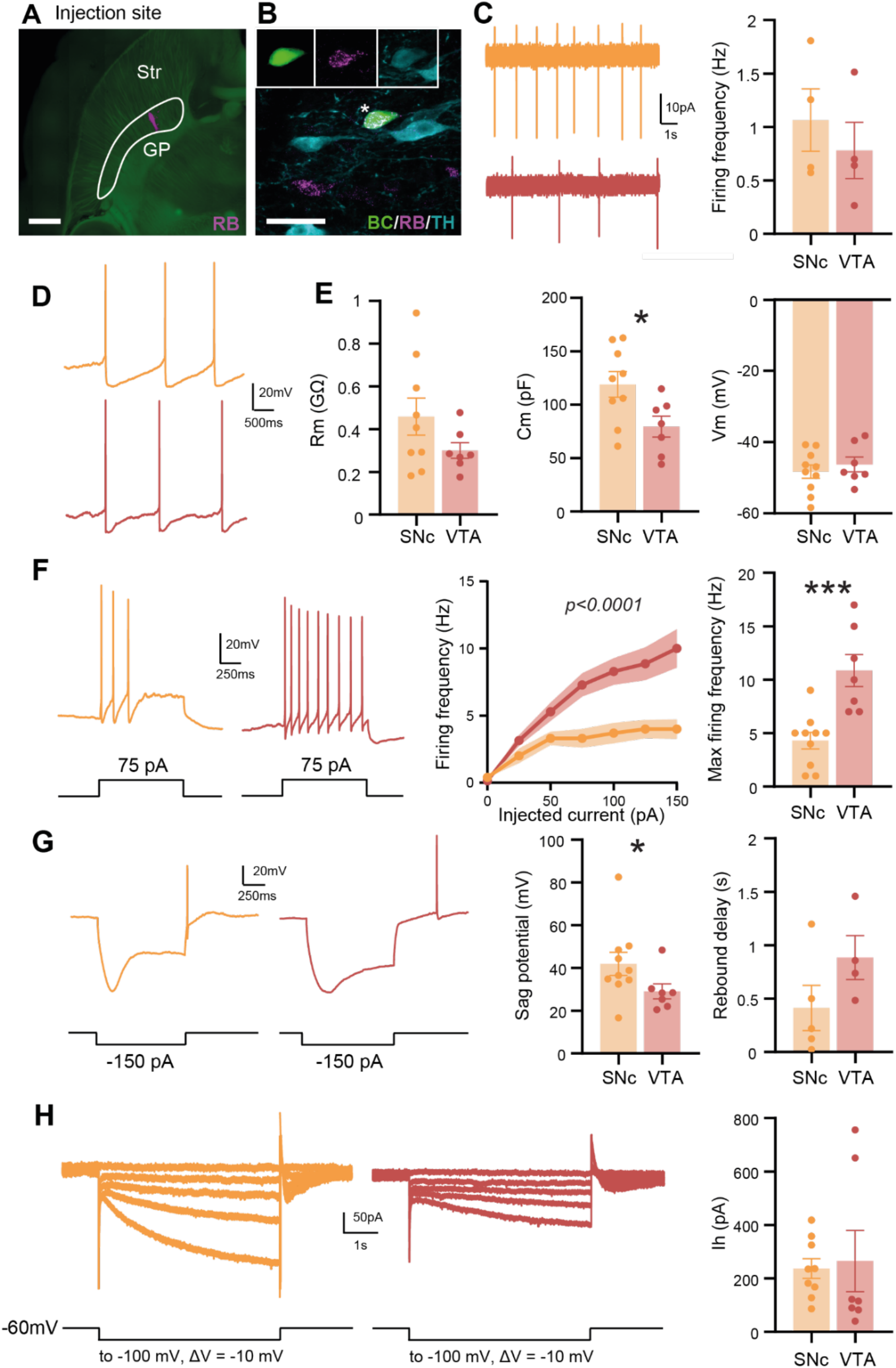
Electrophysiological properties of DA_VTA_ and DA_SNc_ neurons projecting to the GPe. **A**. Representative image showing the injection site of retrobeads (RB) in the GPe. **B**. Example of a recorded neuron filled with biocytin (BC), positive for retrobeads and tyrosine hydroxylase (TH), confirming its identity as a DA neuron projecting to the GPe. **C**. Representative cell-attached recordings from SNc (orange) and VTA (red) neurons and mean spontaneous firing frequency for each population. **D**. Example of whole-cell current-clamp recordings from SNc and VTA neurons. **E**. Summary graphs show membrane resistance (Rm), membrane capacitance (Cm), and resting membrane potential (Vm) for each population. **F**. Representative responses to depolarizing current injections, corresponding F-I curves, and mean maximal firing frequency for each population. **G**. Representative responses to hyperpolarizing current injections in SNc and VTA neurons, and mean sag amplitude and rebound delay for each population. **H**. Representative I_h_ currents from SNc and VTA neurons, generated by 10 mV voltage steps from −60 to −100 mV and mean I_h_ amplitude at −100 mV for each population. Scale bars: 1 mm and 50 μm.

We first verified that retrograde labeling did not affect neuronal viability or intrinsic properties. Membrane resistance (Rm) of labeled neurons was indistinguishable from that of unlabeled

DA neurons, regardless of tracer used (Figure S1; p=0.8785, Kruskal–Wallis), confirming that labeling had no impact on electrophysiology. Because no difference was observed between tracers, data were pooled for subsequent analyses. Retrobeads were preferred for these experiments because they provided stronger labeling and enabled direct visualization of the injection site.

Among neurons exhibiting spontaneous attached-cell activity (47%), mean discharge frequency did not differ significantly between subpopulations (Figure 2C; p=0.4965, unpaired t-test). Passive membrane properties were next examined. While membrane resistance (Rm) and resting potential (Vm) were similar between populations, capacitance (Cm) was significantly higher in DA_SNc→GPe_ compared to DA_VTA→GPe_ neurons (Figure 2E; *p=0.0271, unpaired t-test).

Active properties were assessed by injecting increasing depolarizing currents (Figure 2F). We observed that DA_VTA→GPe_ neurons fired at higher frequencies than DA_SNc→GPe_ neurons (Figure 2F; p<0.0001, Two-Way RM ANOVA) and exhibited a significantly higher maximal firing frequency (Figure 2F; ***p=0.0007, unpaired t-test).

One hallmark of DA neurons is the expression of HCN channels, which generate the hyperpolarization-activated current I_h_. Under current-clamp, HCN-mediated “sag” was evoked by a −150 pA hyperpolarizing step (Figure 2G). Sag kinetics were faster in SNc neurons, and mean sag amplitude was significantly smaller in VTA neurons (Figure 2G; *p=0.0330, Mann– Whitney). Rebound delay was measured in neurons with spontaneous firing. Although not significantly different (p=0.16, unpaired t-test), DA_VTA→GPe_ neurons tended to show longer delays. To directly assess I_h_, we then performed voltage-clamp recordings using hyperpolarizing steps from −60 mV to −100 mV (Figure 2H). Under these conditions, no significant difference in I_h_ amplitude was observed between DA_SNc→GPe_ and DA_VTA→GPe_ neurons. However, substantial variability was noted within the VTA population, with some neurons exhibiting low I_h_ amplitudes and others displaying larger currents.

Together, these data indicate that GPe-projecting DA neurons are electrophysiologically distinct, with marked differences in excitability and sag properties between SNc- and VTA-derived populations.

### The tVTA inhibits DA neurons projecting to GPe

To determine whether DA neurons projecting to the GPe are under inhibitory control from the tVTA, we combined retrograde Fluoro-Gold (FG) tracing from the GPe with anterograde labeling of the tVTA using biotinylated dextran amine (BDA) (Figure 3A-C). We observed that a subset of FG-positive neurons in both the SNc (Figure 3D,d) and the VTA (Figure 3E,e) was closely apposed by BDA-positive fibers originating from the tVTA, suggesting potential synaptic contacts.

**Figure 3.**
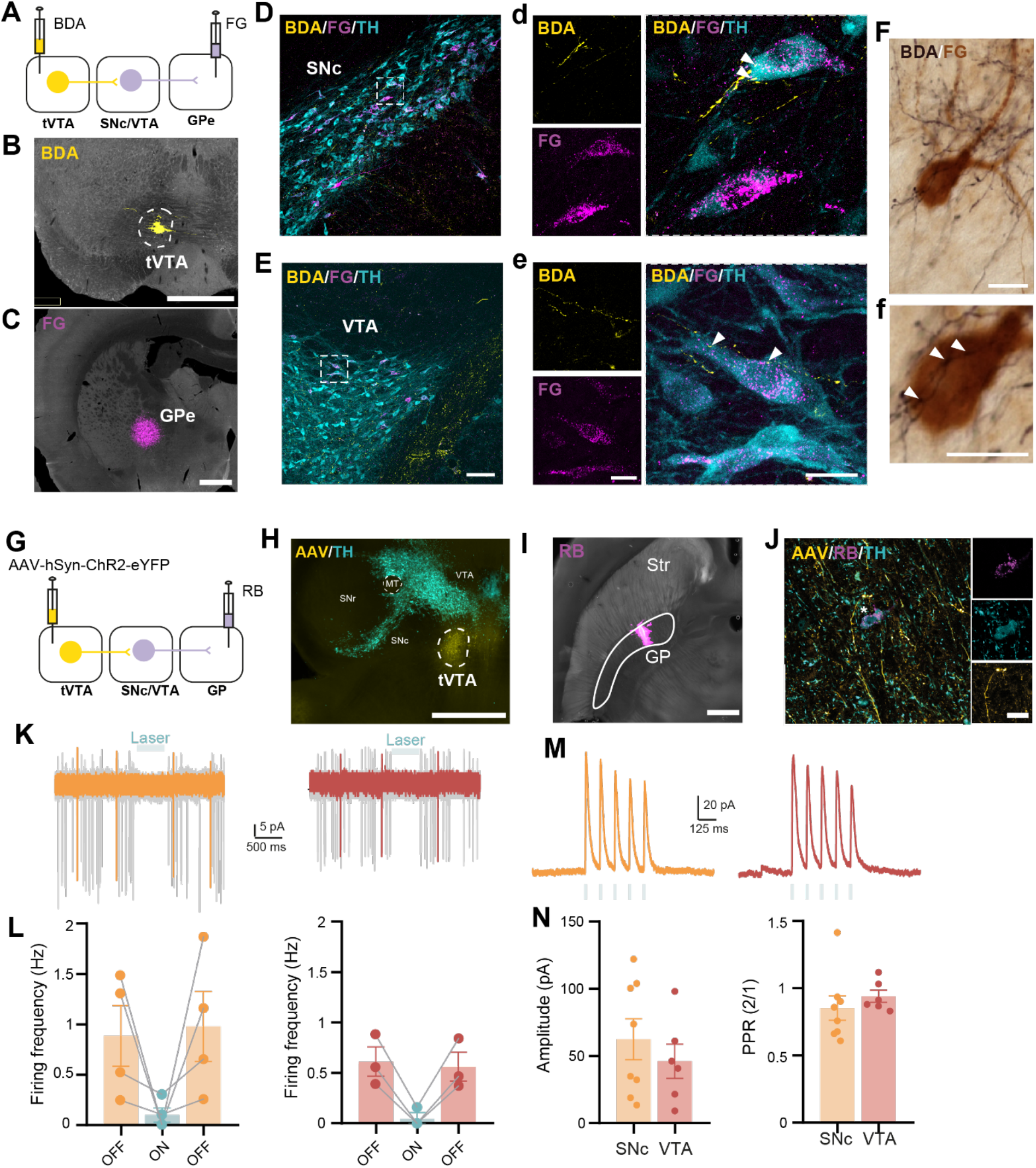
Inhibitory projections from the tVTA onto GPe-projecting VTA and SNc DA neurons. **A**. Schematic of experimental design showing anterograde tracing with biotinylated dextran amine (BDA) injected into the tVTA and retrograde tracing with Fluoro-Gold (FG) injected into the GPe. **B-C**. Representative injection sites of BDA in the tVTA (B) and FG in the GPe (C). **D**. Representative images of the SNc showing TH (blue, DA neurons), FG (magenta, SNc neurons projecting to the GPe), and BDA-labeled fibers (yellow, from the tVTA). **d**. Higher-magnification view showing a SNc DA neuron surrounded by BDA-positive fibers. **E**. Representative images of the VTA showing TH (blue, DA neurons), FG (magenta, VTA neurons projecting to the GPe), and BDA-labeled fibers (yellow, from the tVTA). **e**. Higher-magnification view showing a VTA DA neuron surrounded by BDA-positive fibers. **F**. Immunostaining of BDA (black) and FG (brown) showing a terminal field of tVTA afferents nearby a GP-projecting neuron. **f**. Higher-magnification view showing appositions (arrowheads) of tVTA terminals onto the GP-projecting neuron. **G**. Schematic of optogenetic experiments showing ChR2 expression in the tVTA and retrograde labeling with retrobeads (RB) injected into the GPe. **H-I**. Representative injection sites of AAV-ChR2 in the tVTA (G) and RB in the GPe (H). **J**. Example of a TH^+^ neuron positive for RB and surrounded by AAV-ChR2-positive fibers. **K**. Representative cell-attached recordings from SNc (orange) and VTA (red) DA neurons during optogenetic stimulation of tVTA inputs. **L**. Mean spontaneous firing frequency before, during and after the stimulation for each population. **M**. Representative whole-cell voltage-clamp recordings of light-evoked inhibitory postsynaptic currents (IPSCs) from SNc and VTA DA neurons. **N**. Mean amplitude of IPSCs and paired pulse ratio (PPR) for each population. Scale bars: 1 mm, 100 μm and 20 μm.

To functionally validate this pathway, we performed *ex vivo* electrophysiological recordings combined with optogenetics. A viral vector encoding ChR2-eYFP was injected into the tVTA to enable selective optogenetic stimulation of tVTA neurons (Figure 3G). This approach allowed us to test the existence of functional tVTA projections onto DA_SNc→GPe_ and DA_VTA→GPe_ neurons. In cell-attached configuration, optogenetic stimulation of tVTA neurons induced a robust inhibition of spontaneous firing in both SNc and VTA DA neurons (Figure 3K-L). At the end of the stimulation period, neurons recovered their baseline spontaneous activity. When recorded neurons were silent in cell-attached mode, synaptic responses were assessed in whole-cell voltage-clamp configuration (Figure 3M). Under these conditions, optogenetic stimulation evoked inhibitory postsynaptic currents (IPSCs) that were comparable in DA_SNc→GPe_ and DA_VTA→GPe_ neurons (Figure 3N). Repeated stimulation resulted in synaptic depression, as indicated by a progressive decrease in current amplitude, with no difference in paired-pulse ratio (PPR; 2/1) between the two populations. Bath application of gabazine, a GABA_A_ receptor antagonist, completely abolished the evoked IPSCs, confirming their GABAergic nature. Overall, most of the recorded neurons responded to optogenetic stimulation (88%; 14/16 neurons), either by a decrease in firing frequency in cell-attached mode or by the presence of light-evoked IPSCs in whole-cell recordings.

## DISCUSSION

The present study aimed to identify the anatomical origin of dopaminergic innervation of the GPe and to characterize its control by the tVTA. By combining tracing approaches, immunohistochemistry and *ex vivo* electrophysiology, we demonstrate that dopaminergic inputs to the GPe arise from both the SNc and the VTA and are under strong inhibitory control from the tVTA.

### The GPe receives a dual dopaminergic innervation from SNc and VTA

Using complementary retrograde and anterograde tracing strategies, we demonstrate that, in addition to well-known projections from the SNc (Eid and Parent, 2015; Parent et al., 1990), the GPe also receives substantial dopaminergic input from the VTA. Importantly, the use of selective synaptophysin-based anterograde labeling from SNc and VTA DA neurons serves as a critical control, allowing us to exclude the possibility that tracers injected into the GPe were recaptured by passing fibers.

We next investigated whether these two subpopulations exhibit distinct electrophysiological properties. While some intrinsic properties, such as membrane potential or spontaneous discharge frequency, were similar between the two populations, other parameters revealed clear differences. DA_VTA→GPe_ neurons exhibited higher excitability, as reflected by higher evoked firing frequency. These results are consistent with previous findings showing that VTA DA neurons generally exhibit higher discharge rates than SNc DA neurons under comparable conditions (Lammel et al., 2008).

Previous studies have suggested that DA neurons projecting to the same target share similar electrophysiological features (Ford et al., 2006; Lammel et al., 2008). Our data indicate that DA_SNc→GPe_ closely resemble nigrostriatal neurons, in terms of firing frequency and I_h_ currents (Lammel et al., 2011, 2008). These observations raise the possibility that SNc neurons projecting to the GPe correspond to nigrostriatal neurons sending axon collaterals, as previously suggested by single-neuron reconstruction studies (Matsuda et al., 2009).

In contrast, the DA_VTA→GPe_ population displayed substantial variability, especially in their firing frequency and I_h_ amplitude. Previous work has shown that midbrain DA neurons can be subdivided into two main populations based on these electrophysiological properties. Neurons projecting to the lateral shell of the NAc or to the dorsolateral striatum typically exhibit low firing rates and prominent I_h_ currents, whereas neurons projecting to medial PFC, basolateral amygdala, NAc core and NAc medial shell showed higher maximal discharge frequencies and minimal I_h_ (Lammel et al., 2011, 2008). The variability observed within the DA_VTA→GPe_ population could therefore suggest that these two VTA subpopulations project to the GPe. Future experiments using dual retrograde tracing from the GPe and other target regions will be necessary to determine the extent of overlap between these populations.

The identification of a dual dopaminergic innervation of the GPe raises important questions regarding the respective functional contributions of SNc- and VTA-derived inputs. Although the GPe has long been associated with motor control, it is now increasingly recognized as a complex integrative structure involved in both motor and non-motor processes, including motivation and reward-related behaviors (Beier et al., 2017). Given the established roles of SNc and VTA dopaminergic systems, these observations suggest that SNc-derived inputs may preferentially contribute to motor-related functions of the GPe, whereas VTA-derived inputs could engage circuits related to motivation and learning. Future studies using pathway-specific manipulations will be required to directly test this hypothesis and determine how these distinct inputs shape behavior. Interestingly, the higher excitability observed in DA_VTA→GPe_ neurons suggests that VTA-derived inputs may be more readily recruited than nigropallidal projections. This difference could have important functional implications, potentially biasing GPe activity toward motivational or reward-related signals under certain conditions.

### tVTA exerts strong inhibitory control over GPe-projecting dopaminergic neurons

We show that both DA_SNc→GPe_ and DA_VTA→GPe_ neurons are subject to a strong inhibitory influence from the tVTA. This is in line with previous studies demonstrating dense GABAergic projections from the tVTA onto midbrain DA neurons (Bourdy et al., 2014; Ferreira et al., 2008; Jhou et al., 2009a; Kaufling et al., 2010), as well as its tonic and phasic inhibitory control over their activity (Jalabert et al., 2011; Lecca et al., 2012; Matsui and Williams, 2011; Stamatakis and Stuber, 2012). Our results extend these findings by showing that this regulatory mechanism specifically impacts dopaminergic populations innervating the GPe, suggesting that the tVTA plays a central role in regulating dopaminergic signaling within basal ganglia circuits.

This strong inhibitory control reveals an additional level of regulation within basal ganglia circuits. Given that disorders such as Parkinson’s disease involve the progressive loss of dopaminergic innervation, not only in the striatum but also in the GPe (Benazzouz et al., 2014), modulation of tVTA activity could represent a potential strategy to enhance the function of remaining DA neurons. By relieving tVTA-mediated inhibition, it may be possible to sustain dopaminergic signaling in downstream targets and mitigate motor deficits (Faivre et al., 2020).

Altogether, these findings identify the GPe as a convergent node integrating distinct dopaminergic inputs under tVTA control, suggesting a potential interface between motor and motivational circuits within the basal ganglia.

## RESOURCE AVAILABILITY

Requests for further information and resources should be directed to and will be fulfilled by the lead contact, François Georges

### Materials availability

No new materials were generated in this study.

### Data and code availability

Data and code availability: All information required to reanalyze the data reported in this paper is available from the lead contact upon request.

## ACKNOWLEDGMENTS

This work was supported by the French government through the University of Bordeaux IdEx “Investments for the Future” program (GPR BRAIN_2030). It was also supported by the Centre National de la Recherche Scientifique (CNRS; contract UPR3212), the University of Strasbourg, and the French National Research Agency (ANR; grant ANR-15-CE37-005, tailPARK). We thank Stéphanie Fioramonti for technical assistance. Microscopy experiments were performed at the Bordeaux Imaging Center, a core facility of CNRS, INSERM and the University of Bordeaux, member of the national infrastructure France BioImaging supported by the French National Research Agency (ANR-10-INBS-04). We also thank the Chronobiotron UAR3415 (Strasbourg, France) for animal housing and care, and the In Vitro Imaging platform UAR3156 (Strasbourg, France; France-BioImaging) for microscopy imaging.

## AUTHOR CONTRIBUTIONS

Conceptualization: M.L..; J.B, M.B, and F.G.; Investigation: M.L., G.R.F.; A.B.; Resources: F.G., J.B. and M.B.; Writing – Original Draft: M.L. and F.G.; Visualization: M.L..; Supervision: F.G.; Funding Acquisition: F.G., J.B. and M.B..

## METHODS

### Experimental model and subject details

All experiments were carried out in compliance with the EU Directive 2010/63/EU and the European Committee on Animal Experimentation (CEEA50) under procedures approved by the local ethical committees. These studies were conducted in adult male Sprague-Dawley rats (Charles River, France) and adult male TH-Cre rats (LongEvans-Tg(TH-Cre)3.1Deis). All animals were maintained in a 12/12h light/dark cycle, in stable conditions of temperature and humidity, with access to food and water *ad libitum*.

### Animal surgery

All tracers and viral vectors were microinjected under stereotaxic conditions using glass capillaries (tip diameter: 17-35 μm) connected to a Picopump (Picospritzer III) or to an iontophoresis setup (Midgard, Stoelting, Dublin, Ireland). For surgeries conducted at IMN, animals were anesthetized with isoflurane (induction/maintenance: 5%/1.5%; Centravet, France) in a 50/50 air/O_2_ mixture and placed in a stereotaxic frame (M2E, France). Body temperature was maintained at 37–38°C using a heating blanket (Harvard Apparatus, USA). Ophthalmic ointment (Liposic, Bausch & Lomb) was applied throughout the surgery to prevent eye dehydration. Analgesia was achieved by subcutaneous (s.c.) injection of lidocaine (7 mg/kg) and buprenorphine (0.05 mg/kg). To prevent dehydration, animals received sterile glucose–saline solution (20%, 0.1 mL/10 g body weight, s.c.) immediately after surgery. For surgeries conducted at INCI, animals were anesthetized with an intraperitoneal (i.p.) mixture of Zoletil® 50 (40 mg/kg) and Rompun 2% (10 mg/kg), and placed on a stereotaxic frame (Kopf, Tujinga, CA, USA).

For retrograde tracing experiments, a craniotomy was performed at the following coordinates: AP −1.2 mm, ML 3.4 mm, DV 6 mm, and 30 nL of hydroxystilbamidine methanesulfonate Fluoro-Gold (FG; 2%; Sigma) was injected unilaterally into the GPe through microinjection or iontophoresis (+3 μA with 7s on/off cycles for 7 minutes).

For anterograde tracing experiments, TH-Cre rats received unilateral injections of AAV9-hSyn-Flex-tdTomato-T2A-sypEGFP (Addgene REF:51509) into either the SNc (AP −5.3 mm, ML 2.6 mm, DV 7.5 mm; 180 nL, N=2 rats) or the VTA (AP −5.3 mm, ML 0.7 mm, DV 8.2 mm; 500 nL, N=2 rats).

To visualize tVTA fibers contact on the SNc or on the VTA DA neurons projecting to the GPe, double tracer injections were done by iontophoresis (as described above), with Fluoro-Gold delivered in the GPe and biotinylated dextran amine (BDA, 10000 MW Lysine Fixable, Invitrogen D1956, CA, USA) delivered in the tVTA at the following coordinates, AP: −6.8 mm; ML: 0.4 mm; DV: 7.6 mm (N=4 rats).

For *ex vivo* electrophysiological characterization, retrograde tracers were injected unilaterally into the GPe using similar coordinates: 30 nL of cholera toxin subunit B (CTb-594, 0.5%; Sigma), 30 nL of FG (2%; Sigma), or 60 nL of rhodamine-labeled fluorescent latex microspheres (Lumafluor). To enable optogenetic activation of tVTA neurons, an AAV2-hSyn-ChR2(H134R)-mCherry (UNC; REF: AV4314J) was injected unilaterally into the tVTA at the same coordinates.

### Histology

Rats were euthanized 10–21 days after stereotaxic injections. For IMN-conducted experiments, animals were deeply anesthetized with a mixture of Exagon (200 mg/kg) and lidocaine (20 mg/kg, i.p.) and transcardially perfused with 0.9% NaCl (100 mL), followed by 4% (w/v) paraformaldehyde (PFA) in 0.1 M phosphate-buffered saline (PBS), pH 7.4 (300 mL). Brains were extracted and post-fixed overnight at 4°C in the same fixative solution. The following day, brains were washed twice in PBS and stored in PBS containing 0.03% sodium azide until sectioning. Coronal sections (50 μm) were cut using a vibratome (VT1000S; Leica Microsystems). For the INCI-conducted experiment, rats received an i.p. injection of pentobarbital (364 mg/kg, i.p.; Ceva Santé Animale, Libourne, France) and were transcardially perfused as described above. Following overnight post-fixation, brains were cryoprotected (30% sucrose in Tris-buffered saline, 4°C). 40 μm-thick coronal sections were collected using a freezing microtome (Leica SM2000R microtome).

### Immunohistochemistry

Free-floating sections were washed three times in PBS (10 min each) and then incubated for 1 h at room temperature in PBS containing 0.3% Triton X-100 and 10% normal donkey serum (blocking solution). Sections were incubated overnight at 4°C with primary antibodies diluted in blocking solution: mouse anti-TH (1:1000, Millipore MAB-318), mouse anti-TH (Immunostar #22941, Hudson, WI, USA) or rabbit anti-Fluoro-Gold (Chemicon AB153, Limburg/Lahn, Germany). The following day, sections were washed three times in PBS (10 min each) and incubated for 2 h at room temperature with secondary antibodies diluted in blocking solution. Depending on the experiment, the following secondary antibodies were used: donkey anti-mouse Alexa Fluor 488 (1:500, Invitrogen A-21202), donkey anti-mouse Alexa Fluor 568 (1:500, Life Technologies A-10037), donkey anti-rabbit Cy2 (1:500, Jackson Immunoresearch) or donkey anti-mouse Cy5 (1:500, Jackson Immunoresearch). To visualize BDA, Cy3-conjugated streptavidin was used (1:500 in Tris-HCl, pH 8.2, Jackson Immunoresearch, Ely, UK). Sections were then washed three times in PBS (10 min each) and mounted using antifade mounting medium (Vectashield).

### Imaging and anatomical analyses

Epifluorescence images (AxioImager M2, Zeiss) were acquired using ZEN software (Zeiss) to verify injection sites. Sections containing the SNc and VTA were imaged using a confocal microscope (TCS SPE, Leica or TCS SP5, Leica) and z-stacks acquisitions were performed. Images were processed using ImageJ (NIH) and pseudo colors were applied for visualization. Tracer-labeled neurons in the SNc and VTA were quantified using 5 sections per animal. Cell counting was performed using the Cell Counter plugin in ImageJ.

### *Ex vivo* electrophysiology procedures

Animals were anesthetized with a mixture of ketamine and xylazine (100 mg/kg and 20 mg/kg, respectively) and transcardially perfused with ice-cold sucrose-enriched modified artificial cerebrospinal fluid (sucrose-ACSF), maintained at 95% O_2_ and 5% CO_2_ and containing (in mM): 230 sucrose, 26 NaHCO_3_, 2.5 KCl, 1.25 NaH_2_ PO_4_, 0.5 CaCl_2_, 10 MgSO_4_ and 10 glucose. Brains were removed from the skull, submerged in sucrose-ACSF and cut into 300 um-thick horizontal sections using a vibratome (VT1200S, Leica microsystems). Slices were then immediately transferred in ACSF containing (in mM): 126 NaCl, 2.5 KCl, 1.25 NaH_2_PO_4_·H_2_O, 2 CaCl_2_·H_2_O, 2 MgSO_4_·7H2O, 26 NaHCO_3_, 10 D-glucose, 5 L-glutathion and 1 sodium pyruvate (gassed with 95% O_2_/5% CO_2_). Slices were incubated 1 hour at 32°C in this solution and then maintained at room temperature in the same solution until recording. Sections were then transferred to a recording chamber, continuously perfused with oxygenated ACSF containing (in mM): 126 NaCl, 3 KCl, 1.25 NaH_2_PO_4_·H_2_O, 1.6 CaC_l2_·H_2_O, 1.5 MgSO_4_·7H_2_O, 26 NaHCO_3_, and 10 D-glucose (at 32°C with a perfusion speed of 2 mL.min^-1^). Dopaminergic neurons in the SNc and VTA were visualized using infrared gradient contrast video microscopy (E600FN, Eclipse workstation, Nikon, Japan) and a water-immersion objective (Nikon Fluor 60 X/1.0 NA). Dopaminergic neurons were identified based on morphology and tracer labeling using epifluorescence (Nikon Intensilight C-HGFI). Patch-clamp recordings were obtained using borosilicate glass pipettes (5–8 MΩ) pulled from thick-walled borosilicate glass capillaries (G150–4; Warner Instruments, Hamden, CT, USA) on a micropipette puller (P-97, Sutter Instruments, Novato, CA, USA). The internal solution contained (in mM): 135 K-gluconate, 3.8 NaCl, 1 MgCl_2_·6H_2_O, 10 HEPES, 0.1 Na_4_EGTA, 0.4 Na_2_GTP, 2 MgATP and 5.3 biocytin. The osmolarity and pH of the intrapipette solution were adjusted at 290 mOsm and 7.2 respectively. The recordings were amplified using a Multiclamp 700B amplifier (Molecular Devices) and digitized at 20 kHz (Digidata 1550B) using Clampex 10 acquisition software (Molecular Devices, Sunnyvale, CA, USA).

### Electrophysiological protocols

In cell-attached configuration, spontaneous activity was recorded for 2–4 min at 0 mV. Whole-cell configuration was then established at −60 mV. Passive membrane properties were measured in voltage-clamp mode using −5 mV steps. In current-clamp mode, depolarizing and hyperpolarizing current steps were applied to assess excitability.

Optogenetic stimulation was delivered via a 470 nm LED (Prizmatix) through an optic fiber (500 μm) positioned above the slice. Light pulses (5 ms, 10 Hz) were used to activate tVTA inputs. IPSCs were recorded in voltage-clamp mode and their GABAergic nature confirmed using the GABA_A_ antagonist gabazine (SR95531; 5 μM, Tocris Bioscience).

### Electrophysiology data analysis

Data were analyzed using Clampfit (Molecular Devices) and OriginPro (OriginLab). Membrane resistance (R_m_), capacitance (C_m_) were calculated from responses to −5 mV steps. C_m_ was calculated as Q/ΔV. F-I curves were generated from current-clamp recordings. IPSC amplitudes were measured relative to baseline and paired-pulse ratios were calculated as IPSC_2_/IPSC_1_.

### Cell identification

Following recordings, slices were fixed in 4% PFA and processed for tyrosine hydroxylase (TH) immunostaining. Biocytin-filled neurons were visualized using streptavidin-Alexa Fluor 568 (1:2000, Invitrogen S11226). Fluorescence images were acquired using a confocal microscope (DM5500 TCS SPE, Leica). Z-stack acquisitions with a 2-micron step size were performed using a x40 oil immersion objective. The images were then overlapped using ImageJ (NIH) to confirm the nature of the recorded DA neurons.

### Statistics

Statistical analyses were performed using Prism 8 (GraphPad). Data are presented as mean ± SEM. Normality was assessed prior to statistical testing. Unpaired t-tests or Mann–Whitney tests were used as appropriate. F–I curves were analyzed using two-way ANOVA followed by Sidak’s post hoc test. Data were considered significant for p < 0.05.

