## Supplemental Figure 1 for "tVTA controls dual dopaminergic inputs to the external Globus Pallidus"

### SUPPLEMENTAL INFORMATION

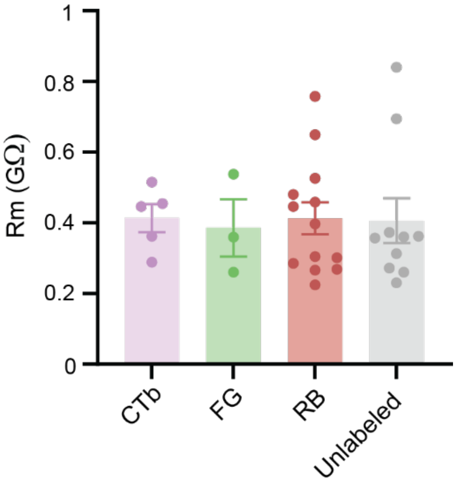


**Figure S1. Membrane resistance of DA neurons is not affected by retrograde labeling.** Membrane resistance was measured in DA neurons that were either unlabeled or labeled with CTb, Fluoro-Gold (FG), or retrobeads (RB). No significant differences were observed between groups, indicating that retrograde labeling does not alter intrinsic electrophysiological properties.
